# Modulation of hippocampal sharp-wave ripples by behavioral states and body movements in head-fixed rodents

**DOI:** 10.1101/2024.12.26.630363

**Authors:** Alain Rios, Usui Minori, Yoshikazu Isomura

**Author notes:** **Correspondence should be addressed to**: Yoshikazu Isomura, PhD, professor, Department of Physiology and Cell Biology, Institute of Science Tokyo, 1-5-45 Yushima, Bunkyo-ku, Tokyo 113-8519, Japan. These authors contributed equally (Alain Rios, Usui Minori).

## Abstract

Hippocampal sharp-wave ripples (SWRs) are critical events implicated in memory consolidation, planning, and the reactivation of recent experiences. Under freely moving conditions, a well-established dichotomy exists: hippocampal networks predominantly generate theta oscillations during periods of reward pursuit (preparatory behaviors) and exhibit pronounced SWR activity once the reward is achieved (consummatory behaviors). Here, we examined how SWRs are modulated by reward delivery and subtle movements in head-fixed rats. Contrary to the canonical view established in freely moving settings, we found that the dominant and more enduring effect was a sustained suppression of SWR activity immediately following water delivery. Moreover, even minor, localized movements (such as whisking or body adjustments) decreased SWR occurrence, demonstrating that hippocampal ripple generation is highly sensitive to motor engagement, irrespective of reward timing. Such movement-induced suppression of ripples persisted during both sleep-like states and quiet wakefulness, suggesting that while large-scale brain states modulate the overall likelihood of SWR generation, local motor-related influences exert a state-independent inhibitory effect on hippocampal ripples. Our results show that SWR modulation by behavioral states and body movements is more context-dependent than previously appreciated.

## Introduction

The hippocampus is integral to learning and memory, functioning within a complex network of brain regions including the prefrontal cortex, medial entorhinal cortex, and frontal cortex (Harvey, 2023). Among the various electrophysiological phenomena observed in the hippocampus, sharp-wave ripples (SWRs) are prominent. These high-frequency, high-amplitude oscillations occur primarily in the CA3 and CA1 regions and are associated with the reactivation of previously active neural ensembles. This reactivation is thought to contribute to memory consolidation, decision-making, planning, and creative thinking (Buzsáki, 2015; Ji & Wilson, 2007). Moreover, disruptions in SWR activity have been implicated in a range of neurological and psychiatric disorders, including epilepsy, schizophrenia, and Alzheimer’s disease (Suh, 2013; Zhen, 2021).

Extensive research has demonstrated that SWRs occur most frequently during states of immobility, sleep, and consummatory behaviors (Buzsáki, 2015; Singer, 2009). Consummatory behaviors fulfill basic physiological needs such as eating, drinking, resting, and defecating (Ball, 2008). Historically, SWRs were thought to primarily facilitate learning and memory consolidation once these needs had been met, supporting the retrospective processing of experiences after immediate drives were satisfied.

However, recent investigations have revealed a more nuanced relationship between SWRs and consummatory behaviors. For instance, recent reports demonstrated a decrease in SWR activity during the intake of water rewards (Klee, 2021), suggesting that these dynamics may be closely modulated by the animal’s current behavioral state. A critical distinction in these studies is whether the animals were head-fixed or allowed to move freely, indicating that SWR patterns can be differentially regulated based on experimental context and the state of motion. These findings underscore the importance of behavioral and environmental factors in shaping hippocampal dynamics.

To further examine how SWRs relate to distinct behavioral states, we conducted experiments in head-fixed rats, simultaneously tracking their movements and recording hippocampal activity. We first compared SWR frequency surrounding reward delivery events (randomly provided water), and subsequently evaluated how changes in the rats’ activity levels related to SWR occurrence throughout the entire session. Through this approach, our study seeks to provide a more comprehensive understanding of the interplay among SWRs, behavioral states, reward processing, and body movements, offering new insights into how hippocampal neural activity aligns with different behavioral states.

## Methods

### Animals and surgery

All experiments were approved by the Animal Care and Use Committee of Institute of Science Tokyo (A2019-274, A2021-041, A2023-116), and were performed in accordance with the Fundamental Guidelines for Proper Conduct of Animal Experiment and Related Activities in Academic Research Institutions (MEXT, Japan) and the Guidelines for Animal Experimentation in Neuroscience (Japan Neuroscience Society). All surgical procedures were performed under appropriate isoflurane anesthesia, and all efforts were made to minimize suffering (Nonomura, 2018; Rios, 2023).

Four wild-type Long-Evans rats (210 ± 35 g, male = 2; female = 2) were kept in their home cages under an inverted light schedule (lights off at 9:00 A.M.; lights on at 9:00 P.M.). The rats were briefly handled by the experimenter (10 min, twice) in advance. For head-plate (CFR-2, Narishige) implantation, the animals were anesthetized with isoflurane (4.5% for induction and 2.0–2.5% for maintenance, Pfizer) using an inhalation anesthesia apparatus (Univentor 400 anesthesia unit, Univentor) and were placed on a stereotaxic frame (SR-10R-HT, Narishige). Lidocaine (Astra Zeneca) was administered around the surgical incisions. Reference and ground electrodes (Teflon-coated silver wires, A-M Systems; 125 μm diameter) were implanted above the cerebellum. During anesthesia, body temperature was maintained at 37 °C using an animal warmer (BWT-100, Bio Research Center). Analgesics and antibiotics were applied postoperatively, as required (meloxicam, 1 mg/kg s.c., Boehringer Ingelheim; gentamicin ointment, 0.1%, MSD). A second surgery was performed for later electrophysiological recordings. We made two tiny holes (1.0–1.5 mm in diameter) in the skull and dura mater above the dorsal hippocampi bilaterally (3.8 mm posterior and ±2.0 mm lateral from the bregma). The skull surface and holes were immediately covered with the antibiotic ointment and silicon sealant until the recording experiments.

After full recovery from surgery (3–7 days later), the rats had *ad libitum* access to water during the weekends, but during the rest of the week they obtained water only during the recording session. Additionally, agar was given to the rats in their home cage to maintain them at >85% of original body weight (Soma, 2017).

### Electrophysiological recording

Extracellular multichannel recordings of local field potentials (LFPs) and single-neuron spike activity were obtained from the hippocampal CA1 area. In two rats, we simultaneously recorded from the parietal cortex located above the CA1 recording sites (approximately 1.2 mm below the cortical surface). Two 32-channel silicon probes (ISO-3x-tet-lin with seven tetrode-like arrangements and four vertically aligned channels on three shanks; NeuroNexus Technologies) were inserted vertically into the left CA1 or the parietal cortex, supported by a 2% agarose-HGT (Nacalai Tesque) gel layer on the brain surface.

Recording depth was determined to detect ripple oscillations within the *stratum pyramidale* (s.p.) of CA1 (Isomura, 2006), typically at the second tetrode from the probe tip. Thus, the upper tetrodes were located in *stratum oriens* (s.o.) and the lower (tip) tetrodes in *stratum radiatum* (s.r.). Probes were inserted at least one hour before the start of each recording session. In the final recording session, probes were coated with the fluorescent dye DiI (DiIC18(3), PromoKine) to visualize probe tracks post hoc.

Neuronal activity was amplified (32-channel main amplifier: FA-32, Multi-Channel Systems; final gain: 2000; band-pass filter: 0.5 Hz to 10 kHz) through a 32-channel head-stage preamplifier (MPA 32I, Multi-Channel Systems; gain: 10). Signals were digitized at 20 kHz using a 32-channel hard-disk recorder (LX-120, TEAC).

### Histology

Following the final recording sessions, rats were deeply anesthetized with urethane (2–3 g/kg, i.p.) and perfused transcardially with cold saline followed by 4% paraformaldehyde in 0.1 M phosphate buffer (PB). The brain was then postfixed and sectioned coronally at 50 µm using a vibratome (VT1000S, Leica, Wetzlar, Germany). Fluorescent DiI-labeled probe tracks and hippocampal CA1 lamination were examined under a fluorescence microscope (BX51N, Olympus).

### SWR and LFP delta power analysis

SWR detection followed standard threshold criteria described previously (Csicsvari et al., 2000; Isomura et al., 2006; O’Neill et al., 2006). Briefly, multichannel signals were down-sampled to 1 kHz and bandpass-filtered in the ripple range (150–250 Hz). The signal envelope was obtained using the Hilbert transform, and SWR events were defined as epochs exceeding 4 standard deviations from the mean (we excluded such events exceeding 9 standard deviations, considering artifacts) for at least 50 ms. The amplitude and peak time of each SWR were defined as the peak amplitude of the enveloped signal in the ripple-band LFP. When SWRs overlapped across multiple channels, the event with the largest peak amplitude was selected for analysis. Artifacts and noise were identified and removed by comparing signals from channels with lower ripple counts and by visual inspection independent of behavior-related information.

For cortical delta power analysis, the LFP signal was bandpass-filtered at 0.5–4 Hz and the envelope was estimated via the Hilbert transform. Periods of delta-on activity were defined as those exceeding 4 standard deviations for at least 200 ms (Girardeau, 2021).

### Video recording and analysis

We used three cameras (Basler ace PoE camera acA1440-73 B/W, Edmund) positioned at front, diagonal, and overhead angles to record the animals’ movements. The initial frame rate (50 frames/s) was frame blended and interpolated to 30 frames/s for analysis using FFmpeg software (FFmpeg team). Movement detection was based on a pixel-wise absolute difference (PWAD) calculation (Soma, 2019):

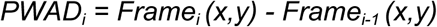

where Frame_i_ represents the *i*th video frame. First, PWAD averages of all frames were obtained for the entire session. Next, regions showing higher levels of movement were identified visually as regions of interest (ROIs), including the orofacial region (tongue, jaw, whiskers, nose), forelimbs, body, and tail. The PWAD values within each ROI were then averaged to generate a continuous pixel-wise difference average (PWDA) per ROI. Movement events were defined as periods where the PWDA exceeded 2.5 standard deviations above the baseline (Musall, 2019).

### Spike analysis

Offline spike sorting was performed to isolate single units from each tetrode. Spike candidates were initially detected and clustered using EToS (Takekawa, 2010; 2012). Manual refinement of these clusters was performed using Klusters and NeuroScope (Hazan, 2006), ensuring that single-neuron clusters had a clear refractory period (>2 ms) in autocorrelograms and no refractory period contamination in cross-correlograms with other clusters. Single-neuron clusters were included if they contained at least 250 spikes in total (Kawabata, 2020; Mitani, 2022).

### Experimental design and statistical analyses

Electrophysiological CA1 data were obtained from 66 recording sessions across 4 rats. In a subset of 12 sessions from 2 rats, we additionally recorded the activity from deep layers of the parietal cortex to evaluate LFP delta power. Each session lasted approximately 90 minutes, during which water (for the sake of simplicity we will call it reward) was delivered through a spout in front of the animal’s mouth at random intervals (180 ± 40 s), with no external cues. Each delivery consisted of three 5-µL drops administered by a micropump. SWR activity was aligned to both water delivery events and ROI-defined movement events to construct peri-event time histograms (PETHs).

SWR frequency was normalized (z-score) on a session-by-session basis using a baseline window (−5 to −2.5 s relative to the event). Two test windows were compared against this baseline: 0 to +0.5 s and 0 to +2.5 s relative to water delivery or movement events. Comparisons were conducted on a trial-by-trial basis within and across sessions using Wilcoxon signed-rank test. Correlations between cortical LFP delta power and CA1 SWR activity were assessed with Pearson’s correlation, and significance was evaluated using a t-test on the correlation coefficients.

All data are expressed as mean ± SD, and n indicates the sample size. Line plots and shaded areas represent mean ± SEM. All statistical analyses were performed with MATLAB’s Statistics and Machine Learning Toolbox (The MathWorks). Differences were considered significant at p < 0.05. Blinding and randomization were not performed.

## Results

### Behavioral States and Movement Quantification during Head-Fixation

To investigate the dynamics of hippocampal sharp-wave ripple (SWR) activity in relation to behavioral states and reward processing, we recorded local field potentials (LFPs) and monitored behavior in head-fixed rats using a three-camera setup. Figure 1 illustrates the overall experimental configuration and the classification of behavioral epochs, as well as representative examples of movement detection within distinct regions of interest (ROIs).

**Figure 1.**
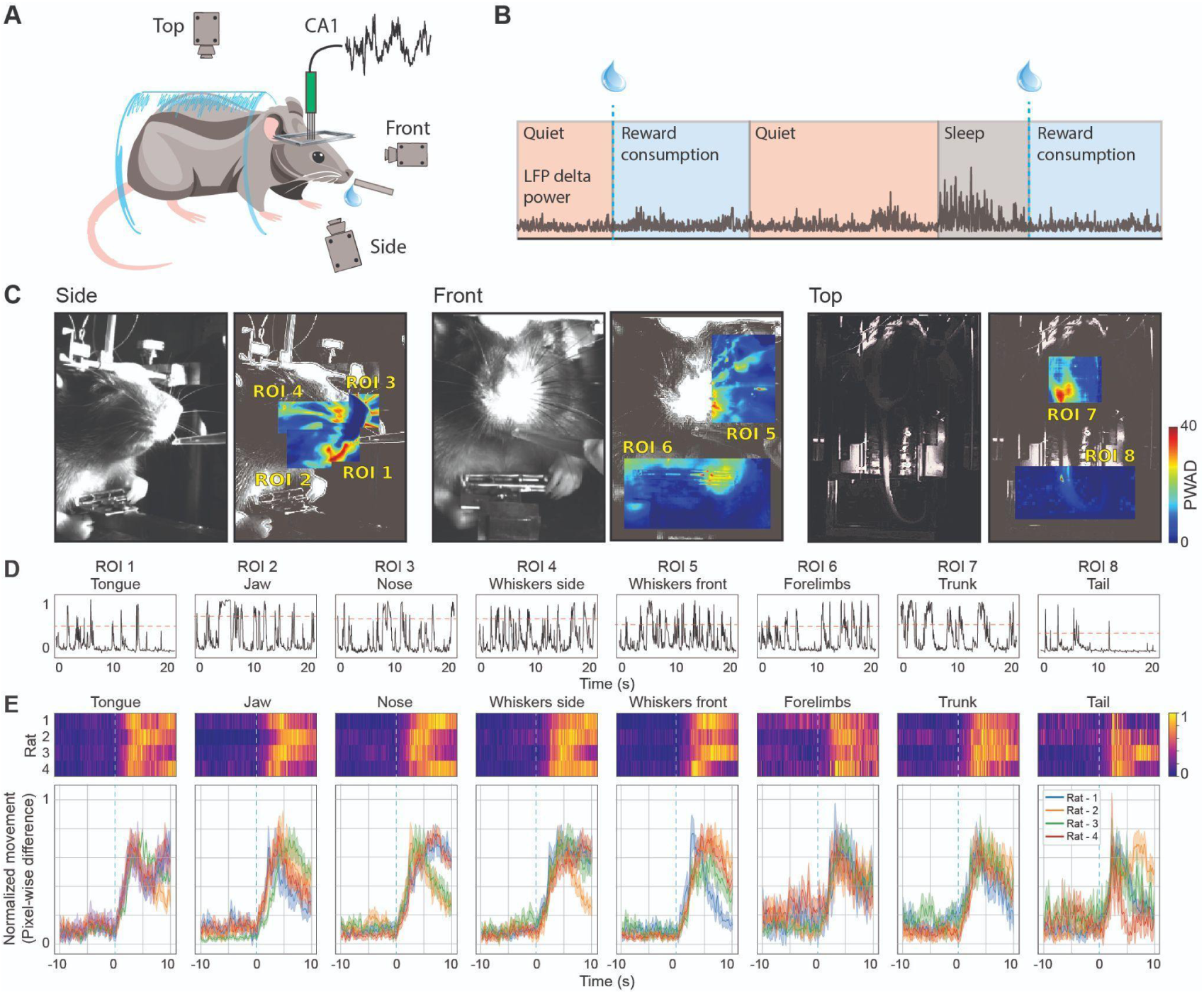
Experimental setup, behavioral state classification, and movement quantification. (**A**) Schematic illustration of the head-fixed experimental configuration. A rat is shown with a recording electrode targeting the hippocampal CA1 region, while three cameras (side, front, top) simultaneously capture the animal’s behavior. Water rewards are delivered through a spout near the rat’s mouth. (**B**) Representative timeline of a recording session segmented into distinct behavioral epochs. Periods of quiet, reward consumption, and sleep are defined based on cortical LFP delta power. The blue drops above the timeline indicate the random delivery of water rewards. (**C**) Example frames from each camera angle (side, front, top) showing the identification of movement ROIs. Left panels: raw video images; right panels: corresponding heat maps illustrating pixel-wise differences highlighting regions with increased movement (ROI 1: tongue, ROI 2: jaw, ROI 3: nose, ROI 4: whiskers side, ROI 5: whiskers front, ROI 6: forelimbs, ROI 7: trunk, ROI 8: tail). (**D**) Representative traces of PWDA of each ROI, showing the threshold used for movement events. (**E**) Top: Colormaps of normalized movement event frequency aligned to reward delivery (time zero) for each ROI, with each row representing a single rat. Bottom: Average perievent time histograms (PETHs) for each ROI, depicting the normalized movement frequency across rats. Shaded areas indicate SEM. Vertical dashed lines mark the time of reward delivery.

As shown in the schematic (Fig. 1A), rats were comfortably head-fixed while we simultaneously recorded hippocampal CA1 LFPs and captured their behavior from three different viewing angles (front, side, and top) with synchronized cameras. The placement of the recording electrode and the camera orientations allowed precise tracking of subtle orofacial and body movements under near-immobile conditions, ensuring that detected movements were not confounded by locomotion.

We segmented the recording sessions into discrete behavioral epochs based on both experimental events and physiological signals. Specifically, reward delivery events (water droplets) were interleaved with intervals of relative immobility. Additionally, we classified distinct states of arousal and behavioral engagement (such as quiet wakefulness, reward-related intervals, and putative sleep episodes) by analyzing cortical LFP delta power (Fig. 1B). Periods with heightened delta power were segregated from those characterized by lower delta and active consummatory behaviors, allowing us to delineate the session into epochs of quiet, reward consumption, and sleep. Quantification of these states (n = 66 sessions, 4 rats) demonstrated consistent transitions between these behavioral epochs across trials.

To assess the animals’ movements in detail, we performed pixel-wise absolute difference (PWAD) analyses on the video frames (see Methods). Representative frames are shown in Figure 1C, with the raw images on the left sub-panels and corresponding ROI-based movement heat maps on the right sub-panels. We defined ROIs for various body parts, including the tongue, jaw, nose, whiskers, forelimbs, trunk, and tail, capturing reward-locked changes in movement. Across all three camera views, these analyses revealed pronounced orofacial and forelimb movements tightly coupled to water reward delivery, as well as more subtle adjustments in body posture and whisking behavior during quiet periods or transitions into rest or sleep states.

Figure 1D provides a quantitative summary of these movement events aligned to reward onset. The top panels show a colormap of normalized movement events for each ROI, averaged across multiple trials. Pronounced increases in orofacial movements were evident shortly before and following reward delivery (tongue: 9.15 ± 1.23 Hz; jaw: 8.45 ± 1.79 Hz; nose: 1.47 ± 0.97 Hz; whiskers side: 1.45 ± 0.82 Hz; whiskers lat: 1.35 ± 0.80 Hz), reflecting the animals’ anticipatory and consummatory responses. In contrast, movements of other body parts (trunk: 0.83 ± 0.54 Hz; tail: 0.71 ± 0.55 Hz) exhibited distinct temporal profiles, often peaking at different times or showing more gradual changes over the trial epoch.

Together, the consistency of movements across animals around reward-related events, and ROI-based analyses provides a detailed behavioral framework against which we will interpret the temporal modulation of SWR activity in subsequent sections.

### Modulation of SWR Activity during consummatory behavior

We next examined how hippocampal SWR frequency and associated CA1 neuronal activity were modulated by reward delivery under head-fixed conditions. To do this, we recorded local field potentials (LFPs) from the CA1 region and aligned SWR events to the timing of water reward administration.

Figure 2A illustrates the analytical framework for detecting SWRs and identifying periods of elevated delta power. In a representative raw LFP trace (top), transient high-frequency oscillations characteristic of SWRs are visible (insets). Filtering the signal in the ripple band (150–250 Hz) and applying the Hilbert transform allowed us to establish a clear threshold (red dashed line) for reliable SWR detection (see Methods). Similarly, applying a low-frequency bandpass (0.5–4 Hz) filter and the Hilbert transform enabled the identification of high-delta epochs. Together, these methods provided a robust and systematic approach for quantifying SWRs and low-frequency activity.

**Figure 2.**
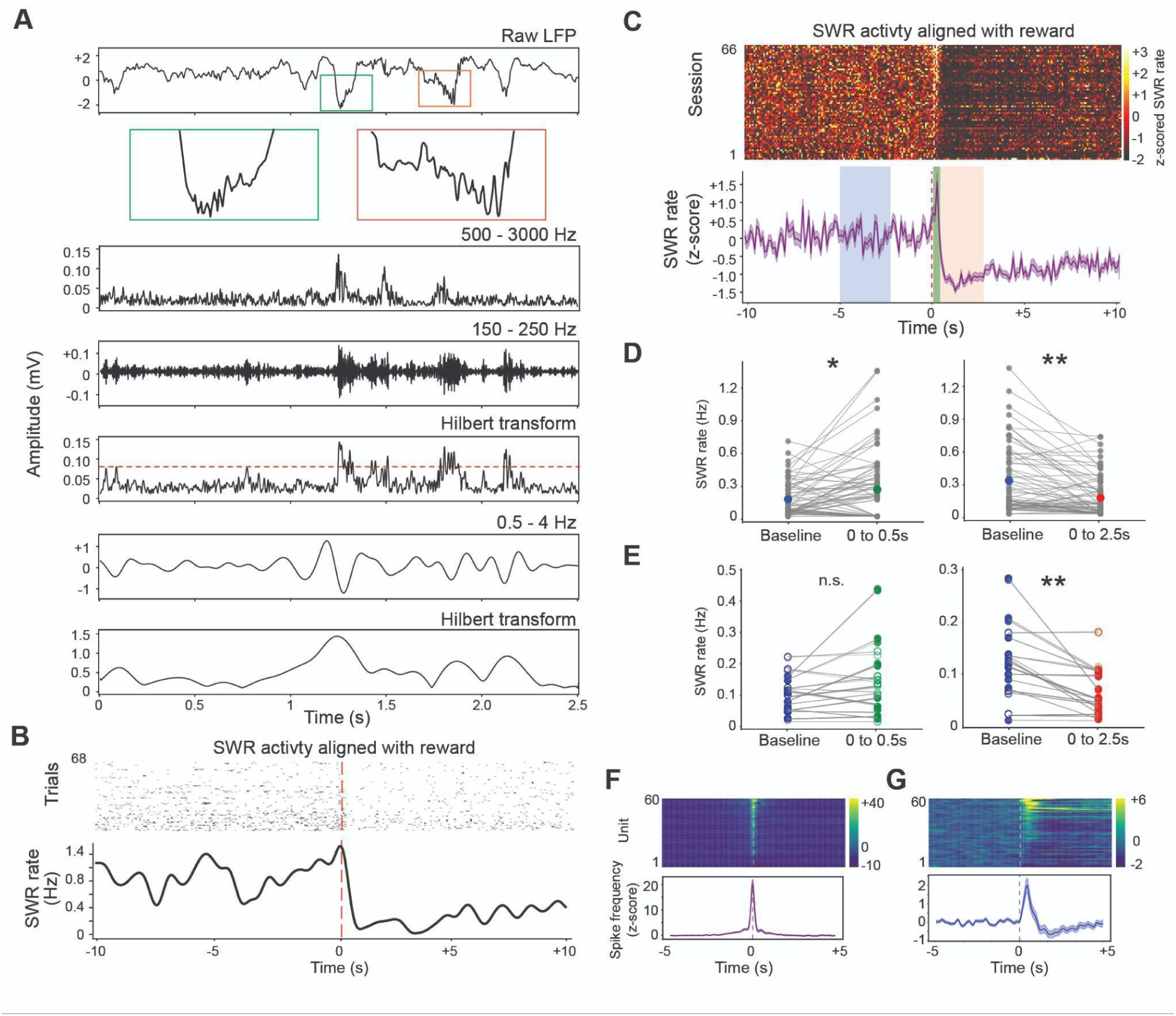
Modulation of hippocampal SWR activity and CA1 spiking by reward events. (**A**) Representative example of SWR and delta-related epochs detection. Top: Raw CA1 LFP trace with two magnified segments (green and red boxes) highlighting transient high-frequency ripple oscillations characteristic of SWRs. Middle: Representative trace of Hilbert transform envelope of bandpass-filtered signal (500–3000 Hz), showing the MUA of the same recording area where SWR were recorded. Below: Bandpass-filtered signal (150–250 Hz) and its Hilbert transform envelope. The red dashed line indicates the SWR detection threshold (4 standard deviations). Further below: Representative 0.5–4 Hz signal and its Hilbert transform envelope used to identify delta-on and delta-off periods. (**B**) Single-session example showing SWR occurrences aligned to reward delivery. Top: Raster plot of SWR events from multiple trials. Bottom: PETH of SWR frequency, demonstrating a phasic decrease in SWR activity immediately following the reward. (**C**) Population-level SWR activity aligned to reward delivery. Top: Colormap of z-scored SWR frequency for all sessions, each row representing one session. Bottom: Mean population PETH, illustrating baseline (blue), short-latency (0–0.5 s, green), and extended (0–2.5 s, red) post-reward intervals. (**D**) Dot plots from a representative session showing trial-by-trial SWR frequency changes from baseline to the short-latency (left) and extended (right) post-reward periods (* = p < 0.05; ** = p < 0.01; Wilcoxon signed-rank test). Color dots indicate mean. (**E**) Population-level comparison of SWR frequency changes, with baseline vs. short-latency (left) and baseline vs. extended (right) post-reward intervals (** = p < 0.01; Wilcoxon signed-rank test). Filled circles denote significant changes (either increases or decreases), whereas empty circles represent non-significant sessions. (**F**) Cross-correlograms of single-unit spiking relative to SWR maxima for a subset of sessions (top: all recorded neurons as rows; bottom: average cross-correlogram). (**G**) Population spike firing rates of CA1 neurons around reward delivery.

Next, we evaluated how SWRs were modulated around the time of reward delivery. As shown in Figure 2B, a representative session revealed a pronounced decrease in SWR frequency immediately following reward administration. This trial-by-trial visualization demonstrates that the reward onset reliably influenced SWR timing, leading to a significant reduction relative to the pre-reward baseline.

We extended this analysis to the entire dataset (Figure 2C, see Table 1 for summary of statistical findings). In some sessions we observed a brief increase of SWR activity around reward delivery. Two post-reward intervals were tested: a short-latency window (0–0.5 s, green) and a more extended interval (0–2.5 s, red). While some sessions exhibited a transient, phasic increase in SWR frequency immediately after reward delivery (Figure 2D; p = 0.018 for 0–0.5 s; p = 1.52e-4 for 0–2.5 s; Wilcoxon signed-rank test), the population-level analysis revealed that the predominant effect was a prolonged decrease in SWR activity following the initial response (Figure 2E; p = 2.01e-5, Wilcoxon signed-rank test for 0–2.5 s after reward). On the other hand, the transient increase in SWR frequency observed in certain sessions was not consistent across the population. Thus, although the reward could induce short-lived elevations in ripple frequency in some cases, the dominant and consistent pattern across sessions was a sustained suppression of SWRs over the longer post-reward interval.

**Table 1.**

| Figure | Data | Rats | Primary statistic | Comparison | Statistic |
| --- | --- | --- | --- | --- | --- |
| 2D | SWR rate (Hz) | N=1 rat | Wilcoxon signed-rank test | Baseline vs 0-0.5s window | n = 59 trials, W = 510, p = 0.018 |
| 2D | SWR rate (Hz) | N=1 rat | Wilcoxon signed-rank test | Baseline vs 0-2.5s window | n = 59 trials, W = 730, p = 1.52e-4 |
| 2E | SWR rate (Hz) | N=4 rats | Wilcoxon signed-rank test | Baseline vs 0-0.5s window | n = 66 sessions, W = 213, p = 0.24 |
| 2E | SWR rate (Hz) | N=4 rats | Wilcoxon signed-rank test | Baseline vs 0-2.5s window | n = 66 sessions, W = 945, p = 2.01e-5 |
| 3A | SWR rate (Hz) | N=4 rats | Wilcoxon signed-rank test | Baseline vs 0-2.5s window | n = 66 sessions, W = 809, p = 1.22e-03 |
| 3B | SWR rate (Hz) | N=4 rats | Wilcoxon signed-rank test | Baseline vs 0-2.5s window | n = 66 sessions, W = 998, p = 4.78e-02 |
| 3C | SWR rate (Hz) | N=4 rats | Wilcoxon signed-rank test | Baseline vs 0-2.5s window | n = 66 sessions, W = 845, p = 4.50e-02 |
| 3D | SWR rate (Hz) | N=4 rats | Wilcoxon signed-rank test | Baseline vs 0-2.5s window | n = 66 sessions, W = 556, p = 4.83e-02 |
| 3E | SWR rate (Hz) | N=4 rats | Wilcoxon signed-rank test | Baseline vs 0-2.5s window | n = 66 sessions, W = 735, p = 7.44e-03 |
| 3F | SWR rate (Hz) | N=4 rats | Wilcoxon signed-rank test | Baseline vs 0-2.5s window | n = 66 sessions, W = 691, p = 3.22e-03 |
| 3G | SWR rate (Hz) | N=4 rats | Wilcoxon signed-rank test | Baseline vs 0-2.5s window | n = 66 sessions, W = 905, p = 4.47e-05 |
| 3H | SWR rate (Hz) | N=4 rats | Wilcoxon signed-rank test | Baseline vs 0-2.5s window | n = 66 sessions, W = 845, p = 4.50e-02 |
| 4B | SWR band, Delta band | N=1 rat | Pearson's correlation | SWR band power, Delta band power | n = 60 trials, $r^2 = 0.65$; p = 0.0002 |
| 4C | SWR band, Delta band | N=2 rat | T-test for correlation | SWR band power, Delta band power | mean Pearson's correlation coeff: $0.54 \pm 0.14$, p = 6.53e-08 |
| 4D | SWR rate (Hz) | N=2 rat | Wilcoxon signed-rank test | Baseline vs 0-2.5s window. Delta on epochs. | ROI-1: n = 12 sessions, W = 34, p = 1.61e-04;<br>ROI-2: n = 12 sessions, W = 18, p = 0.0181;<br>ROI-3: n = 12 sessions, W = 31, p = 0.0018;<br>ROI-4: n = 12 sessions, W = 8, p = 0.0593;<br>ROI-5: n = 12 sessions, W = 29, p = 1.18e-03;<br>ROI-6: n = 12 sessions, W = 46, p = 2.57e-09;<br>ROI-7: n = 12 sessions, W = 52, p = 1.71e-12;<br>ROI-8: n = 12 sessions, W = 48, p = 2.59e-10 |
| 4E | SWR rate (Hz) | N=2 rat | Wilcoxon signed-rank test | Baseline vs 0-2.5s window. Delta off epochs. | ROI-1: n = 12 sessions, W = 2, p = 0.56933;<br>ROI-2: n = 12 sessions, W = 20, p = 0.0203;<br>ROI-3: n = 12 sessions, W = 31, p = 0.00172;<br>ROI-4: n = 12 sessions, W = 16, p = 0.0771;<br>ROI-5: n = 12 sessions, W = 20, p = 0.02757;<br>ROI-6: n = 12 sessions, W = 22, p = 0.00016;<br>ROI-7: n = 12 sessions, W = 10, p = 0.0321;<br>ROI-8: n = 12 sessions, W = 3, p = 0.1920 |

To validate our SWR detection approach and assess the functional relevance of these events, we examined their relationship with neuronal firing (Adaikkan, 2024). We observed an increase in the Multi-unit activity (MAU) related to a high ripple band power (Figure 1A). Additionally, using a subset of recordings (n = 12 sessions) where single-unit activity was simultaneously acquired, we constructed cross-correlograms between neuronal firing times and SWR peaks (Figure 2F). This analysis revealed a significant increase in spike discharge coincident with SWR events, confirming that the detected ripples corresponded to genuine hippocampal network activity.

Finally, we examined changes in spike firing rates of CA1 neurons around reward delivery (Figure 2G). Consistent with the notion that reward information can transiently enhance hippocampal excitability (Singer, 2009), most neurons showed increased firing shortly after reward. Nonetheless, a subset of neurons exhibited no significant change or even a reduction in firing, highlighting the heterogeneity of hippocampal network responses to reward events.

These results demonstrate that reward delivery under head-fixed conditions modulates hippocampal SWR activity, with a complex temporal pattern characterized by transient increases in some sessions but a more robust, long-lasting suppression observed across the population.

### Modulation of SWRs by Orofacial and Body Movements

We next investigated the relationship between SWR occurrence and the onset of movement events associated with different body regions. To avoid confounding influences of reward-related effects on ripple activity, we excluded all movement events occurring within ±12 s of reward delivery. With this constraint in place, we aligned SWR activity to the onset of detected movements in specific ROIs, including the tongue, jaw, nose, whiskers (lateral and front), forelimbs, trunk, and tail. We then compared SWR frequencies in a baseline window preceding the movement event to a test window immediately following it.

Orofacial movements led to a predominant decrease in SWR frequency across sessions (Fig. 3A–E). Although a minority of sessions showed increased or unchanged SWR activity, the overall trend was a significant reduction relative to baseline levels. This reduction was similarly observed for movements of other body parts, including forelimbs, trunk, and tail (Fig. 3F–H). Although session-to-session variability existed (with some sessions showing increases or no change), the dominant population-level pattern across all these ROIs was a reduction in SWR activity associated with movement events. In each case, the population PETHs revealed a downward shift in SWR frequency following the onset of the corresponding body-part movement, with the session-based dot plots confirming a significant proportion of sessions exhibiting this downward modulation.

**Figure 3.**
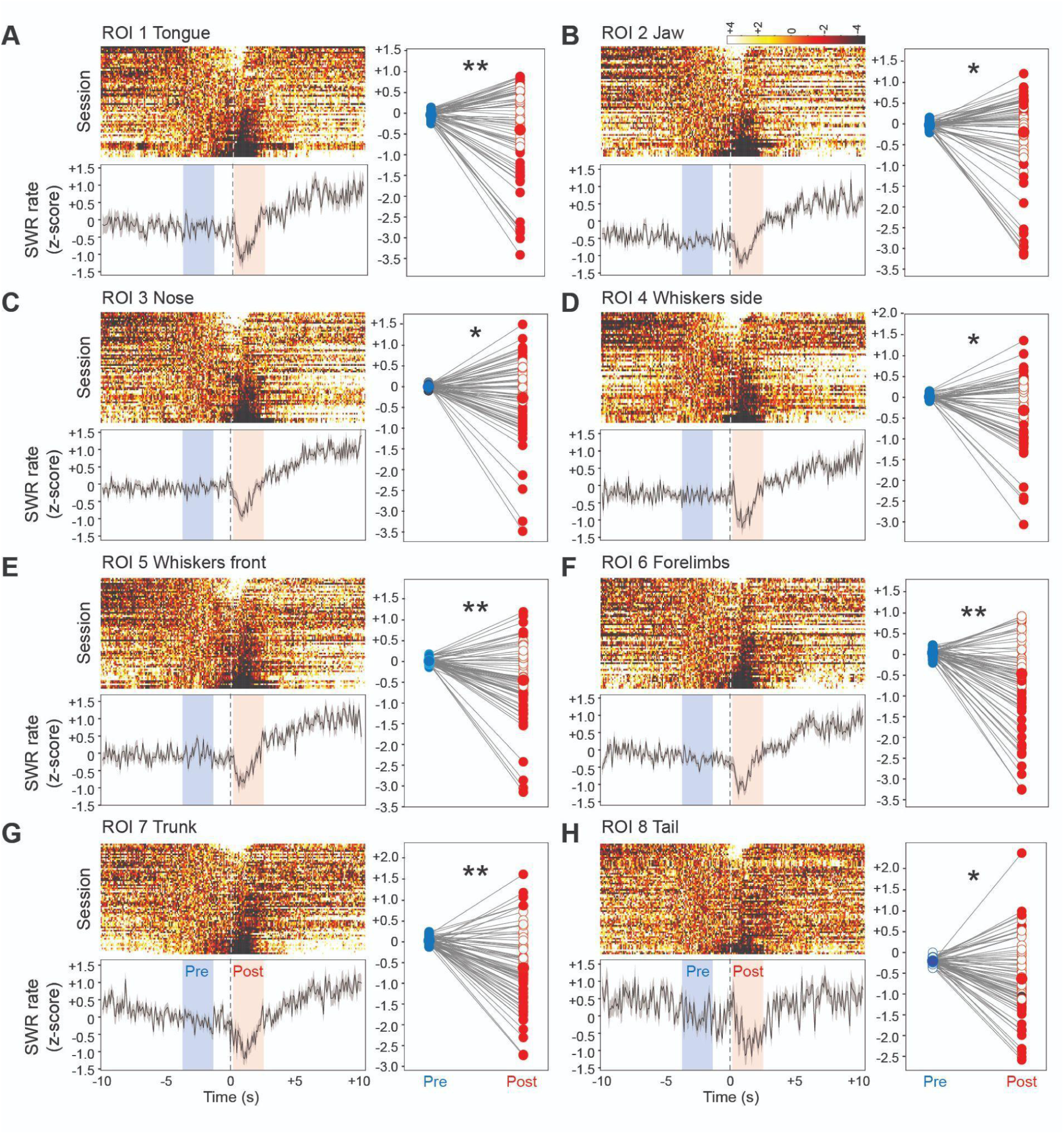
Modulation of SWR activity by orofacial and body movements, independent of reward proximity. (**A**) Left top: Population-level heatmap of SWR activity aligned to the onset of tongue movements. Each row represents one session, with color indicating changes in SWR frequency from baseline. While a subset of sessions showed increases or no significant change, the majority exhibited a decrease in SWR frequency. Left bottom: Population averaged PETH of SWR activity centered on tongue higher movement displacement. Baseline (blue) and test (light red) windows are indicated, revealing an overall reduction in SWR activity following the movement. Right: Dot plot comparing baseline and test windows on a per-session basis, with filled symbols indicating sessions that reached statistical significance. (**B**) SWR activity aligned to jaw movements. (**C**) SWR activity aligned to nose movements. (**D**) SWR activity aligned to whisker lateral movements. (**E**) SWR activity aligned to whisker front movements. (**F**) SWR activity aligned to forelimb movements. (**G**) SWR activity aligned to trunk movements. (**H**) SWR activity aligned to tail movements. (* = p < 0.05; ** = p < 0.01; Wilcoxon signed-rank test).

Together, these findings indicate that hippocampal SWRs are generally suppressed around the time of active orofacial and body movements, even when excluding movements that occur near reward delivery. This suggests that hippocampal network states conducive to SWR generation are less prominent when the animal engages in even minor, localized movements. This relationship was consistent across multiple ROIs, underscoring the robust link between behavioral activity and hippocampal ripple dynamics.

## Delta Power Correlates with SWR Band Activity and Influences Movement-Related Modulations

To further explore the interplay between behavioral states and SWRs, we examined how fluctuations in cortical delta power (0.5–4 Hz) correlate with hippocampal ripple activity (150–250 Hz).

Figure 4A presents a representative example of simultaneously recorded cortical delta-band and CA1 SWR-band filtered LFP signals, both enveloped and normalized for clarity. Visual inspection revealed that elevations in delta power frequently coincided with increases in SWR activity. This synchronous modulation aligns with previous studies linking large-scale brain states, such as sleep or quiescent wakefulness, to coordinated changes in hippocampal ripple generation (Goldstein, 2018; Mölle, 2006; Sirota, 2003).

**Figure 4.**
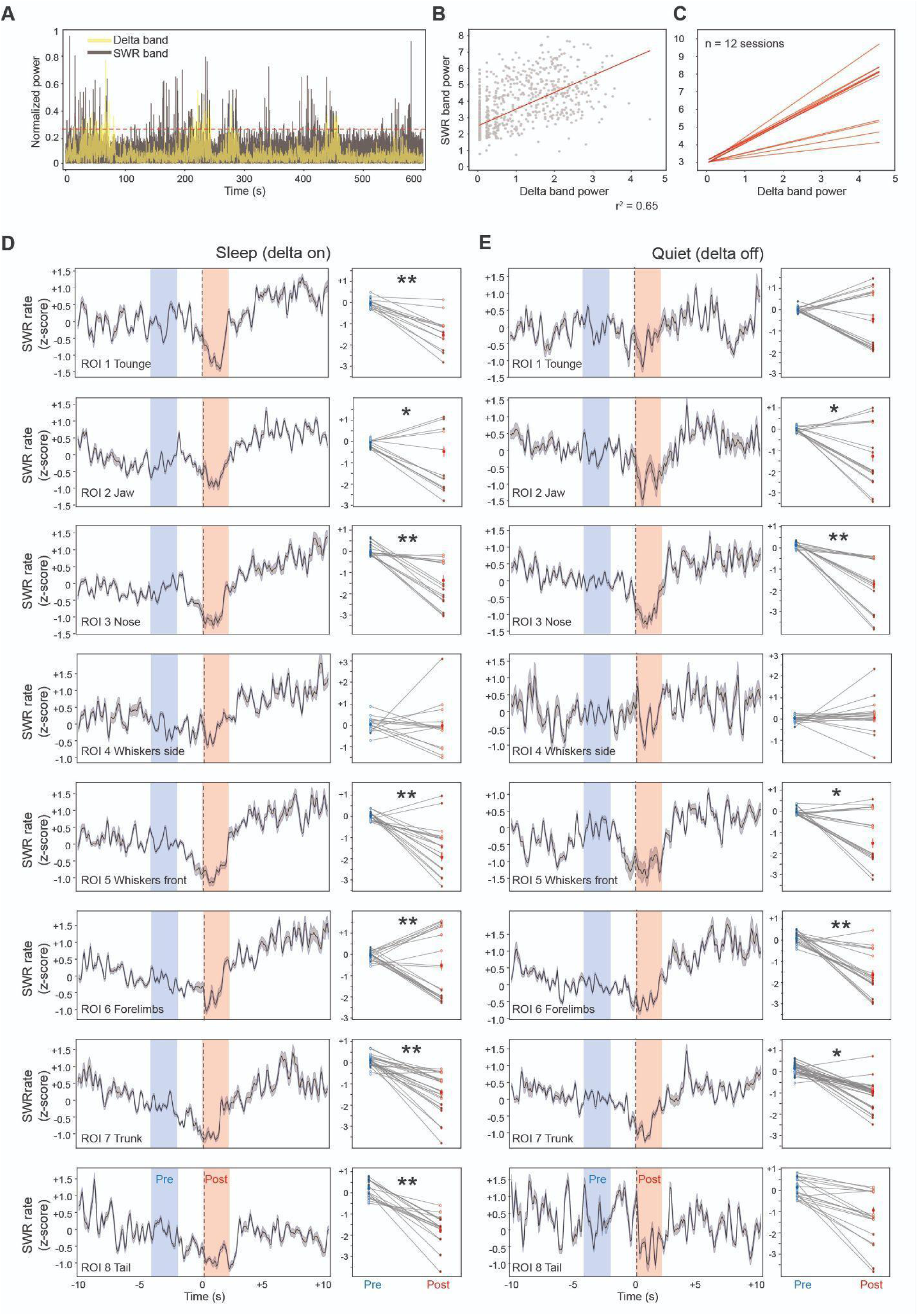
Relationship between cortical delta power and hippocampal SWRs, and its impact on movement-related SWR modulation across behavioral states. (**A**) Representative example of simultaneously recorded delta-band (0.5–4 Hz, yellow) and SWR-band (150–250 Hz, gray) power traces from cortical and CA1 LFPs, respectively. Both signals are enveloped and normalized. (**B**) Scatter plot of SWR-band power versus delta-band power from a single representative session. Each point corresponds to a time bin (200 ms), and the red line indicates a best-fit linear regression. A significant positive correlation (r² = 0.65; p = 0.0002) demonstrates that increases in delta-band power are accompanied by concurrent increases in SWR activity. (**C**) Correlation analysis of delta and SWR power extended across all recorded sessions (n = 12). Each line represents a single session’s regression line. A one-sample t-test of the Pearson correlations confirms that the positive relationship between delta and SWR power is robust and consistent across the population (mean Pearson correlation: 0.54 ± 0.14, p = 6.53e-08). (**D**) Movement-related modulation of SWRs during delta-on (sleep-like) epochs. For each ROI (tongue, jaw, nose, whiskers lateral, whiskers front, forelimbs, trunk, tail), the left panels show the average PETHs of SWR frequency aligned to movement events. Blue and red shaded areas represent the baseline and test windows, respectively. The right panels illustrate session-by-session comparisons of pre- vs. post-movement SWR activity (blue and red dots, respectively), with filled symbols indicating sessions that reached statistical significance. (**E**) The same analysis as in (D), but during delta-off (quiet awake) epochs. (* = p < 0.05; ** = p < 0.01; Wilcoxon signed-rank test).

To quantify this relationship, we performed a correlation analysis between delta-band and SWR-band power within individual sessions. As shown in Figure 4B, a single session exhibited a significant positive correlation (r² = 0.65, p = 0.0002), indicating that shifts in delta power were accompanied by concurrent changes in SWR band activity. Extending this analysis across all recorded sessions (n = 12), we found that the positive correlation between delta and SWR power was consistent and robust (Figure 4C). On average, delta-band power reliably tracked fluctuations in SWR activity (mean Pearson correlation: 0.54 ± 0.14, p = 6.53e-08, one-sample t-test), reinforcing the notion that delta oscillations and ripples are dynamically coupled and that large-scale cortical states influence hippocampal network excitability.

Given the clear link between delta power and SWR occurrence, we next sought to disentangle the effects of brain state (i.e., delta-on as a putative sleep-like state vs. delta-off as an awake, quiet state) on SWR modulation around movement events. We defined delta-on epochs as those exceeding four standard deviations above the mean delta envelope amplitude, interpreting these periods as sleep-like. Conversely, delta-off epochs were considered awake, quiescent states.

In Figure 4D, we focused on the delta-on (sleep-like) epochs and repeated the movement-related SWR analysis (as in Figure 3) for each ROI (tongue, jaw, nose, whiskers lateral, whiskers front, forelimbs, trunk, and tail). During delta-on states, the onset of movement generally coincided with a decrease in SWR frequency, though the degree of modulation sometimes varied across sessions. These findings indicate that although SWRs are coupled to delta power, the movement-related suppression of SWRs persists during sleep-like epochs.

During delta-off (quiet awake) epochs, we again observed that movement onset typically reduced SWR frequency, despite a higher number of sessions displaying non-significant changes or increases compared to sleep-like states. While the overarching pattern of movement-induced SWR suppression remained consistent across delta-on and delta-off conditions, subtle differences in the magnitude or significance of SWR modulation suggest that the underlying brain state, as reflected by delta power, can influence the extent to which hippocampal ripples are affected by behavioral events.

In summary, these results demonstrate that SWR activity is tightly coupled to cortical delta oscillations and that this relationship remains evident when examining the effects of movement on SWRs across different brain states. Whether during delta-on (sleep-like) or delta-off (awake quiet) epochs, movement-related suppression of SWRs persists, underscoring the multifaceted and state-dependent regulation of hippocampal network dynamics.

## Discussion

In this study, we investigated how hippocampal SWRs are modulated by behavioral states, reward delivery, and localized movements in head-fixed rats. Building on previous work that established SWRs as critical for memory consolidation, decision-making, and planning (Ji & Wilson, 2007; Buzsáki, 2015; Girardeau, 2011) and that their disruption can lead to pathological outcomes (Suh et al., 2013; Zhen et al., 2021), our findings provide new insights into the dynamic interplay between hippocampal network activity, motivational factors, and subtle behavioral fluctuations.

Traditionally, research conducted in freely moving rodents has outlined a clear dichotomy between preparatory and consummatory behavioral states (Buzsáki, 2015; O’Keefe & Nadel, 1978; Csicsvari et al., 2007). During preparatory periods, when animals actively pursue rewards, hippocampal activity is dominated by theta oscillations, reflecting heightened spatial and attentional processing. In contrast, when animals transition into a consummatory phase (such as consuming a reward) SWRs typically increase, suggesting the facilitation of the consolidation and replay of recently acquired information (Buzsáki, 2015; Singer & Frank, 2009). This distinction is well-documented under freely moving conditions, where animals engage in naturalistic sequences of foraging, reward pursuit, and consumption.

Our results challenge and refine this mainstream understanding. Under head-fixed conditions, we observed a far more complex and context-dependent modulation of SWRs by reward delivery. While some sessions displayed a transient, short-lived increase in SWR frequency immediately following reward (Fig. 2), the predominant pattern was a prolonged suppression of SWRs across the population (Fig. 5). The majority of significant change across sessions and body parts is decrease in SWR activity, further corroborating the notion that even subtle movements suppress hippocampal ripple events in a broad range of behavioral contexts. In a similar direction, during delta-off epochs, regardless of whether some sessions and ROIs display no significant changes or occasionally an increase in SWRs, the dominant pattern remains a suppression of SWR activity around movement events. This finding is consistent with recent observations reporting similar results in head-fixed rats (Klee, 2023). Thus, although hippocampal networks retain the capacity for SWR enhancement under certain conditions, head fixation and the associated constraints on naturalistic movement and behavioral sequences appear to reshape the interplay between reward consumption and ripple generation. The lack of full-bodied locomotion and extensive exploratory behaviors in head-fixed paradigms may reduce or alter the mnemonic demands and spatial contexts that drive the classic preparatory-consummatory dichotomy observed in freely moving animals.

**Figure 5.**
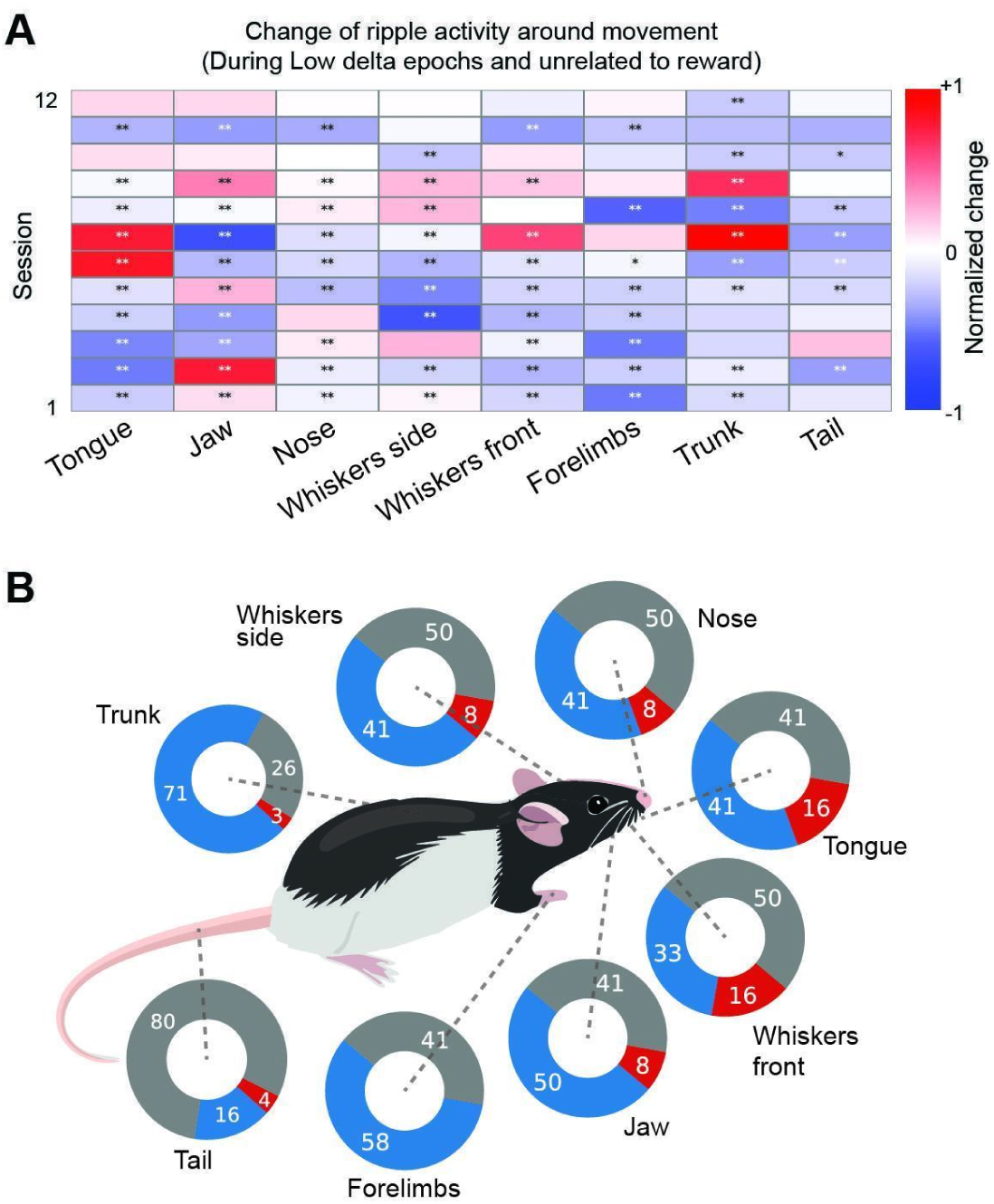
Comprehensive summary of movement-related changes in hippocampal SWR activity during delta-off, reward-free epochs. (**A**) Session-by-session matrix illustrating changes in SWR frequency aligned to the onset of movements for multiple body parts during delta-off epochs (tongue, jaw, nose, whiskers side, whiskers front, forelimbs, trunk, tail). Each row represents one recording session, and each column corresponds to a specific body part. Colors indicate the normalized change in SWR activity relative to baseline: blue denotes a decrease, and red denotes an increase. Statistical significance, assessed by Wilcoxon signed-rank test, is indicated by asterisks (*p < 0.05; **p < 0.01). (**B**) Schematic summary of the SWR activity related to body movements. Each body part of the rat is paired with a donut plot indicating the proportion of sessions that showed significant decreases (blue), significant increases (red), or no significant change (gray) in SWR activity.

Beyond reward modulation, we found that even minor orofacial and body movements strongly suppressed SWR occurrence (Figs. 3–4). This phenomenon extends the classic view that SWRs occur predominantly during immobility, quiet wakefulness, and sleep (Buzsáki, 2015; Foster & Wilson, 2006). Our results reveal that subtle, localized movements, far from the vigorous locomotion typically studied, are sufficient to disrupt the conditions favoring SWR generation. While in freely moving rodents, preparatory states often involve exploratory locomotion that can suppress ripples (Kay et al., 2019; Sullivan et al., 2011), our findings suggest that even in a head-fixed setting, minimal motor engagement still reallocates hippocampal resources away from internal replay and toward processing immediate sensory-motor inputs. Thus, the suppression of SWRs during subtle movements may represent a fundamental and conserved principle of hippocampal function, one that transcends differences in the environmental complexity or degree of physical freedom available to the animal.

Our analyses also showed that SWR activity is closely coupled with cortical delta oscillations (Fig. 4), a known hallmark of global brain states such as sleep and quiet wakefulness (Mölle et al., 2006; Sirota et al., 2003). Whether in delta-on (sleep-like) or delta-off (quiet awake) epochs, the fundamental movement-related suppression of SWRs persisted (Fig. 5B). This finding implies that while delta power modulates the overall excitability landscape for SWR generation, it does not override the inhibitory influence of movement. The interaction between global brain states and local motor-related signals allows the hippocampus to flexibly balance mnemonic processing with the demands of the current environment (Jadhav et al., 2012; Ólafsdottir et al., 2018).

Our results further underscore the complexity of SWR regulation: reward and consummation, which freely moving studies have often associated with enhanced SWR activity (Singer & Frank, 2009), here yielded a more context-sensitive pattern. The restricted and highly controlled conditions of head fixation likely reduce spatial navigational demands and the variety of preparatory actions that characteristically precede reward consumption in free-ranging foraging scenarios. As a result, hippocampal circuits may shift away from replaying reward-related experiences and instead maintain a lower excitability state after a brief post-reward transient. This difference highlights the importance of behavioral context and environmental constraints in interpreting hippocampal activity patterns and their functional significance.

Though our head-fixed preparation allowed precise monitoring of subtle movements and provided stable electrophysiological recordings, it also limited the behavioral repertoire and spatial exploration of the animals. Future research should replicate these experiments in freely moving settings to confirm the generality of our findings, particularly regarding the dissociation between preparatory and consummatory states (for example, under more cognitive demanding tasks, requiring a higher engagement in attention, discrimination or decision making). Additionally, exploring how SWR modulation by movement and reward differs across hippocampal subregions or species could clarify the universality of these regulatory mechanisms.

In summary, our study provides an alternative view of SWRs as dynamic phenomena influenced not only by global brain states and reward conditions but also by subtle, localized movements (Fig. 5C). By comparing our results with the classical paradigms of freely moving animals, we highlight that SWR modulation is highly context-dependent. This contributes to a more nuanced understanding of hippocampal function, demonstrating that even within ostensibly similar behavioral states (e.g., consummatory behavior), the neural underpinnings of SWR generation can shift depending on the sensory-motor context and environmental constraints. Such flexibility ensures that the hippocampus can continuously adapt its memory-related activity amidst the rich and ever-changing tapestry of experience.

## Conflict of interest

The authors declare no competing financial interests.

## Acknowledgments

This work was supported by Grants-in-Aid for Scientific Research (B) (JP19H03342 and JP23H02589 to Y.I.), for Transformative Research Areas (A) (JP21H05242 to Y.I.), and for Challenging Research (Exploratory) (JP24K21999 to Y.I.) from MEXT and JSPS; by the Takeda Science Foundation (Y.I.); and by Center for Brain Integration Research, Institute of Science Tokyo. We appreciate all members of Isomura laboratory, especially Drs. Masanori Kawabata, Riichiro Hira, and Yutaka Sakai.

## Notes

### Competing Interest Statement

The authors have declared no competing interest.

